# Comprehensive identification of potentially functional genes for transposon mobility in the *C. elegans* genome

**DOI:** 10.1101/2023.08.08.552548

**Authors:** Yukinobu Arata, Peter Jurica, Nicholas Parrish, Yasushi Sako

**Affiliations:** Cellular Informatics Laboratory, Cluster for Pioneering Research (CPR), RIKEN, 2-1 Hirosawa, Wako, Saitama, 351-0198, Japan; Genome Immunobiology RIKEN Hakubi Research Team, RIKEN Center for Integrative Medical Sciences, 1-7-22 Suehiro-cho, Tsurumi-ku, Yokohama City, Kanagawa, 230-0045, Japan

## Abstract

Transposons are mobile DNA elements that encode genes for their own mobility. Whereas transposon copies accumulate on the genome during evolution, many lose their mobile activity due to mutations. Here, we focus on transposon-encoded genes that are directly involved in the replication, excision, and integration of transposon DNA, which we refer to as “transposon-mobility genes”, in the *Caenorhabditis elegans* genome. Among the 62,773 copies of retro- and DNA transposons in the latest assembly of the *C. elegans* genome (VC2010), we found that the complete open reading frame structure was conserved in 290 transposon-mobility genes. Critical amino acids at the catalytic core were conserved in only 145 of these 290 genes. Thus, in contrast to the huge number of transposon copies in the genome, only a limited number of transposons are autonomously mobile. We conclude that the comprehensive identification of potentially functional transposon-mobility genes in all transposon orders of a single species can provide a basis of molecular analysis for revealing the developmental, aging, and evolutionary roles of transposons.

## Introduction

Transposon-related repetitive elements occupy a significant proportion of eukaryotic genomes: 12% in *Caenorhabditis elegans*, 4-9% in *Drosophila*, 37% in mouse, 46% in human, and ~85% in corn (Huang et al., 2012). During evolution, transposons increase their copy number via transposition but lose their mobile activity due to mutations, insertions, and deletions within the transposon (Huang et al., 2012; Kidwell and Lisch, 2001). Depending on the transcript required for transposition, transposons are primarily classified into retrotransposons (class I) and DNA transposons (class II) (Kapitonov and Jurka, 2008; Wells and Feschotte, 2020; Wicker et al., 2007). Each class is further subdivided into orders by the transposition mechanism. The 5 retrotransposon orders are the long terminal repeat (LTR), long interspersed nuclear element (LINE), *Dictyostelium* intermediate repeat sequence (DIRS), Penelope-like element (PLE), and short interspersed nuclear element (SINE) orders. The 4 DNA transposon orders include the terminal inverted repeat (TIR), Helitron, Maverick/Polinton (MP), and Crypton orders (Wells and Feschotte, 2020).

The LTRs transpose by replicating their DNA copies from the transcript by reverse transcriptase (RT). Integrase (IN) inserts free transposon DNA fragments into other genomic loci. Similarly, LINEs transpose by replicating their DNA copies from the transcript by LINE-specific RT. In this case, reverse transcription starts at the 3’-end of the nick site on the genomic DNA generated via the endonuclease (EN) domain at the N-terminus of the RT. DIRS transposition occurs when a free circular double-stranded DNA is generated via reverse transcription. The circular double-stranded DNA is then integrated into another locus by Tyrosine-Recombinase (Tyr-REC) (Cappello et al., 1985; Goodwin and Poulter, 2001, 2004; Poulter and Butler, 2015). In PLE transposition, the DNA fragment of PLE is reverse transcribed from the 3’ end of the nick site on the genome generated by the GIY-YIG EN activity in the PLE-specific RT (Pyatkov et al., 2004). The TIR DNA transposons transpose via the EN activity of transposase (TP). The TP recognizes terminal inverted repeats at both ends of TIR and cleaves double-stranded DNA (dsDNA) to generate a free transposon DNA fragment. This fragment is integrated into another genomic locus by the EN activity of TP (Hickman and Dyda, 2016; Hickman et al., 2005; Nesmelova and Hackett, 2010). Helitron transposition occurs when the Helitron DNA element is nicked via the EN activity of the REP domain in REP-Helicase (REP-HEL) and is then unwound into single-stranded DNA (ssDNA) through HEL activity. Although it remains controversial whether the transposon ssDNA is replicated before integration into another genomic locus, a free DNA fragment is integrated into another locus by EN activity of the REP domain (Chandler et al., 2013; Grabundzija et al., 2016; Kapitonov and Jurka, 2001, 2007; Thomas and Pritham, 2015). In MP DNA transposons, free DNA copies are replicated by DNA Polymerase B encoded within MP. These DNA copies are inserted into other genomic loci by MP-encoded INs (Feschotte and Pritham, 2005, 2007; Gao and Voytas, 2005; Kapitonov and Jurka, 2006). Cryptons transpose after being excised as free circular DNAs (circDNAs) by Tyr-REC. The free circDNAs are integrated into another genome locus by Tyr-REC (Goodwin et al., 2003; Poulter and Butler, 2015).

Thus, all orders of transposons are transposed via 2 processes: 1) the production of a free transposon DNA, and 2) its integration into another genomic locus. In retrotransposons LTR and DIRS, and DNA transposon MP, the free transposon DNA is generated and integrated by different enzymes. In retrotransposons LINE and PLE, and DNA transposon Helitron, these processes are mediated by different enzymatic activities associated with each functional domain in the same enzyme. In DNA transposons TIR and Crypton, these processes are mediated by the same functional domains in the same enzyme. Hereafter, we refer to the genes involved in the production and/or integration of free transposon DNA copies as “transposon-mobility genes”.

The retrotransposon L1 is a member of the well-studied LINE order of transposons. The human genome has more than 500,000 copies of L1, corresponding to 17% of the human genome (Cordaux and Batzer, 2009). However, bioinformatic analysis has identified only 146 copies of full-length L1 in the human genome, and only 107 of these L1 copies have conserved intact RT genes (Penzkofer et al., 2017). Experimental tests of the transposon mobility of L1 showed that fewer than 100 copies of L1 were active, with 6 L1 copies accounting for most of the activity of L1 in the human genome (Brouha et al., 2003). These results raise the possibility that, in contrast to the huge number of transposon copies in the genome, only a few transposons in other orders are autonomously mobile. Nevertheless, to date, no research has comprehensively explored how many transposons among all orders in a single species are autonomously mobile.

Here, we focus on transposon-encoded genes that are directly involved in the replication, excision, and integration of transposon DNA, which we refer to as “transposon-mobility genes”, in the *C. elegans* genome. We bioinformatically searched the latest *C. elegans* genome assembly (VC2010) for transposon-mobility genes that conserve the complete open reading frame (ORF) structure and critical amino acids at the catalytic core for enzymatic activities. Our analysis uncovered most of the transposon-mobility genes that were identified in previous studies (Bessereau, 2006; Bowen and McDonald, 1999; Fischer et al., 2003; Ganko et al., 2001; Kanzaki et al., 2018; Kapitonov and Jurka, 2001; Marín et al., 1998; Youngman et al., 1996), as well as novel genes.

## Results

### Comprehensive identification of transposon-mobility genes in the *C. elegans* genome

To identify transposon copies on the latest *C. elegans* genome assembly (VC2010), we used RepeatMasker (RM), an algorithm that identifies repetitive DNA elements in a genome (https://www.repeatmasker.org/). Our RM analysis showed that repeat elements occupy 13.86% of the *C. elegans* genome (Table 1). This result is consistent with previous reports that 12-17% of the *C. elegans* genome is repeat elements (Kanzaki et al., 2018; Laricchia et al., 2017; Stein et al., 2003). After excluding simple, satellite, and low-complexity repeat elements, we identified 62,773 copies of transposons. To identify protein-coding elements among these 62,773 transposon copies, we used an *ab initio* gene finder, SNAP, which was shown to identify 97% of ORFs on the *C. elegans* genome (Korf, 2004). Our SNAP analysis identified 808 potential gene-coding regions. Among them, 428 genes conserved the complete ORF structure. To infer the function of genes encoded in these complete ORFs, we used DIAMOND (Double Index AlignMent Of Next-generation sequencing Data), an algorithm for fast and sensitive protein alignment (Buchfink et al., 2014). DIAMOND identified 285 transposon-mobility genes (80 RT, 11 IN, 189 TP, and 5 REP-HEL genes).

**Table 1:**
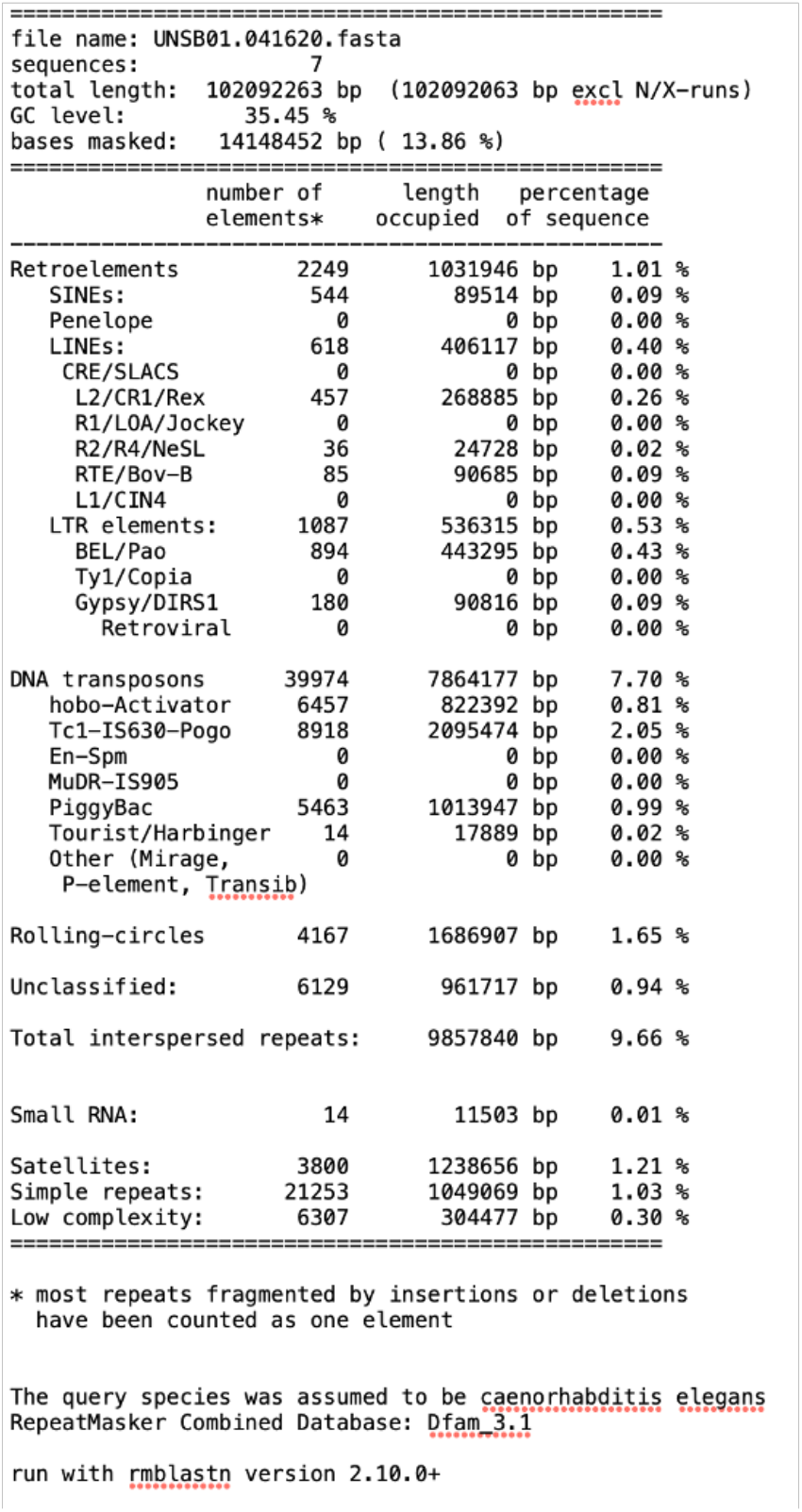

### Transposon-mobility genes in LTRs in *C. elegans*

Among the 80 RT genes, 19 genes were encoded in LTRs (*rtz_LTR*s) and 61 genes were encoded in LINEs (*rtz_LINE*s) (Supplementary Tables 1 and 3). RTs, *i.e.* RNA-dependent DNA polymerases, have 5 evolutionarily conserved motifs (A, B’, C, D, and E) (Poch et al., 1989; Xiong and Eickbush, 1990). Asp residues in Motifs A and C are widely conserved in all RNA/DNA-dependent DNA polymerases and DNA/RNA-dependent RNA polymerases (Delarue et al., 1990). Previous crystal structure analyses showed that one Asp in Motif A and 2 Asp residues in Motif C form a catalytic triad for holding a bivalent metal ion for conjugating the alpha phosphate of a new dNTP to the OH group to the 3’ end of the DNA strand (Le Grice, 2012; Jacobo-Molina et al., 1993; Steitz, 1998). Mutations in Asp residues of Motifs A or C reportedly abolish RT activity (Larder et al., 1987). By aligning RTZ_LTRs with the reference RTs, we identified 8 RTZ_LTRs that conserved Asp residues in Motifs A and C (red asterisks in Figure 1A). Additionally, these 8 RTZ_LTRs preserved 1) the conserved Gly residue in Motif B’ (Smith et al., 2006), which functions for interacting with the incoming nucleotide and template strand (Mitchell et al., 2010), and 2) the Leu and Gly residues in Motif E, which function for fixing the primer strand and positions it toward the active site (Mitchell et al., 2010) (Figure 1C).

**Figure 1.**
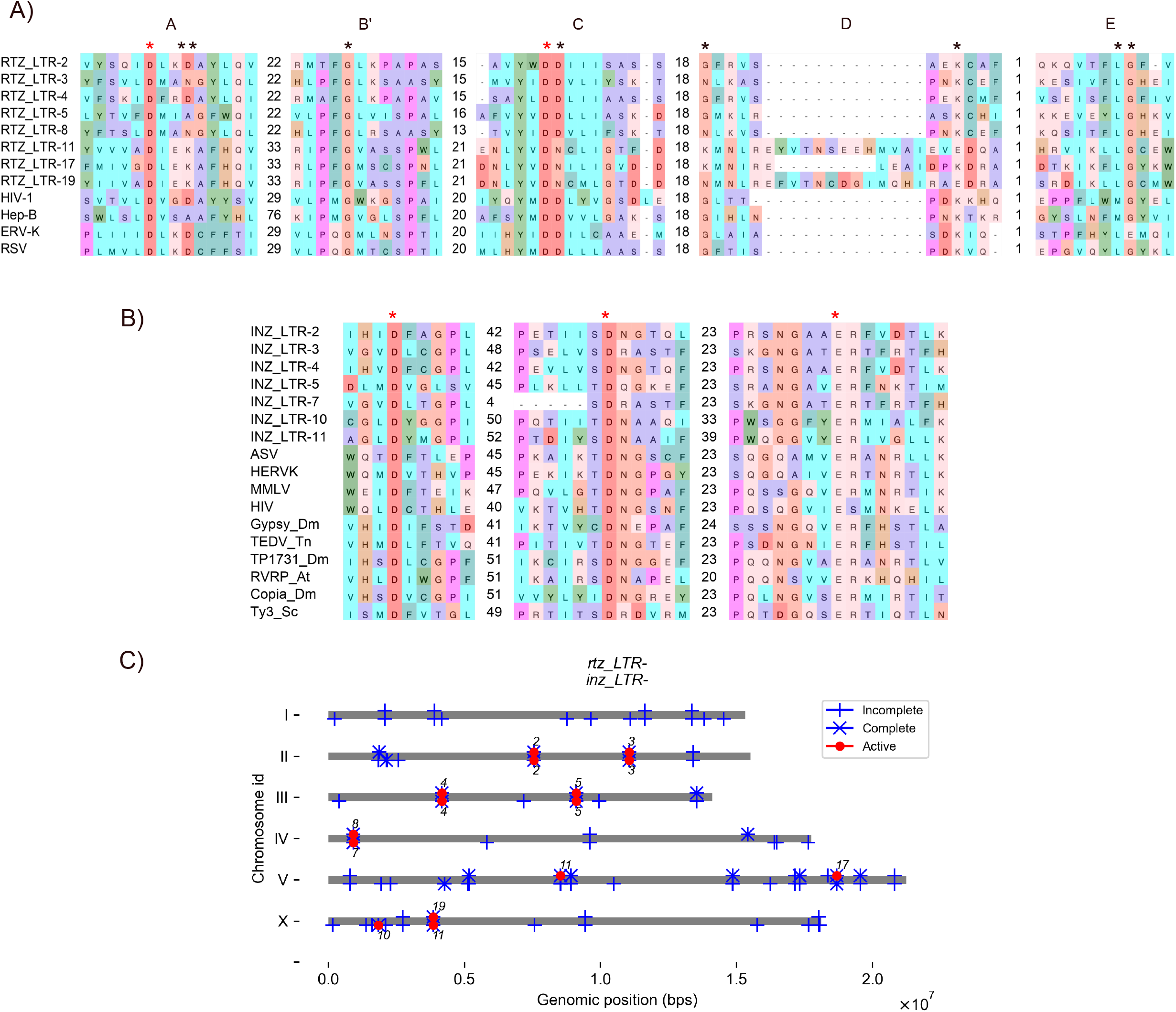
Potentially functional *rtz_LTR*s and *inz_LTR*s encoded in long terminal repeats (LTRs) (A) Alignment of catalytic core domains of RTZ_LTRs with reference reverse transcriptases (RTs). Abbreviations: HIV-1: Chain C, HIV-1 RT P66 subunit of human immunodeficiency virus type 1 [5TXO_C], Hep-B: Hepatitis B virus RT_like family [QFR04538], ERV-K: Pol protein of human endogenous retrovirus K [CAA76882], RSV: Pol of Rous sarcoma virus [CAA48535]. (B) Alignment of catalytic core domains of INZ_LTRs with reference integrases (INs). Abbreviations: ASV: IN in avian sarcoma virus [1ASU_A], HERVK: Pol protein in human endogenous retrovirus K [CAA76885], MMLV: p46 IN in Moloney murine leukemia virus (MoMLV) [NP_955592.1], HIV: IN in human immunodeficiency virus type 1 [1BIZ_A], Gypsy_Dm: IN, Gypsy endogenous retrovirus in *D. melanogaster* [CAB69645], TEDV_Tm: ORFB in TED virus in *T. ni* [YP_009507248], TP1731: Pol polyprotein in transposon_1731 in *D. melanogaster* [S00954], RVRP_At: retrovirus-related like polyprotein in *A. thaliana* [CAB78488_1], Copia_Dm: Gag-Int-Pol protein in COPIA in *D. melanogaster* [P04146], Ty-3_Sc: Gag-Pol polyprotein in Ty3-G in *S. cerevisiae* [GFP69998.1]. Asterisks indicate conserved residues. Red asterisks indicate residues for identifying potentially functional transposon-mobility genes. (C) Genomic positions of 8 potentially functional *rtz_LTR*s and 7 potentially functional *inz_LTR*s in *C. elegans* genome. Grey lines represent chromosomes. Red circles over and under chromosome indicate positions of potentially functional genes of *rtz_LTR*-n and *inz_LTR*-n, respectively. Numbers over and under chromosome indicate numbers of *rtz_LTR* and *inz_LTR* genes, respectively. Vertical ticks and × marks with vertical ticks over and under chromosome indicate incomplete and complete ORFs, respectively, of *rtz_LTR*s and *inz_LTR*s, respectively.

In RTZ_LTR-11, -17, and -19, the second Asp residue in Motif C was substituted with Asn (black asterisk in Figure 1A). As described above, the second Asp residue is involved in a catalytic triad. Site-directed mutagenesis at the second Asp residue, including mutagenesis to Asn as found in RTZ_LTR-11, -17, and -19, significantly reduces RT activity in human immunodeficiency virus (HIV) (Boyer et al., 1992; Le Grice et al., 1991; Kaushik et al., 1996). However, amino acid substitution at the second Asp to Asn has been found in functional RNA-dependent RNA polymerases in negative-strand RNA viruses (Barik et al., 1990; Delarue et al., 1990; Poch et al., 1989). Therefore, we considered the possibility that RTZ_LTR-11, -17, and -19 might still be functional.

In RTZ_LTR-11 and -19, the conserved Gly and Lys in Motif D were substituted. In RTZ_LTR-3, -8, and -17, the conserved Gly in Motif D was substituted. Amino acid substitutions at Gly in Motif D are often observed in RNA-dependent RNA polymerases in negative-strand RNA viruses (Poch et al., 1989). Therefore, we considered the possibility that RTZ_LTR-3, -8, and -17 might still be functional. On the other hand, Lys in Motif D is highly conserved throughout polymerases (Poch et al., 1989; Xiong and Eickbush, 1990). Motif D functions for forming a phosphodiester bond with dNTPs with the 3’-OH (hydroxyl) of the primer (Canard et al., 1999; Castro et al., 2009). Motif D in RTZ_LTR-11 and -19 may be nonfunctional. Nevertheless, given the conservation of the 2 critical amino acids holding a bivalent metal ion, and the fact that we did not perform functional testing, we held out the possibility that RTZ_LTR-11 and -19 might be functional. Taken together, these results led us to classify all 8 *rtz_LTR*s as potentially functional genes (Supplementary Table 1).

Next, we identified 11 IN genes encoded in LTRs (*inz_LTR*s) (Supplementary Table 2). The catalytic core domain of IN has an evolutionarily conserved DD35E motif that is required for EN activity (Engelman and Craigie, 1992; Van Gent et al., 1992; Kulkosky et al., 1992). DD35E holds 2 metal ions, which integrate a free transposon DNA into the genome (Hare et al., 2012; Maertens et al., 2021). Amino acid substitution at the 3 critical amino acid residues in the DD35E motif abolishes EN activity of INs in Rous sarcoma virus (RSV) and HIV (Drelich et al., 1992; Engelman and Craigie, 1992; Van Gent et al., 1992; Kulkosky et al., 1992). By aligning the 11 INZ_LTRs with the reference INs, we identified 7 INZ_LTRs that conserved the DD35E triads (Figure 1B and Supplementary Table 2). Therefore, we considered these 7 *inz_LTR*s as potentially functional genes. All of the *inz_LTR*s except *inz_LTR-10* were encoded in the same LTR with the potentially functional *rtz_LTR* genes (Figure 1C).

### Transposon-mobility genes in LINEs in *C. elegans*

By aligning the 61 RTZ_LINEs with reference RTs, we identified 28 RTZ_LINEs that had Asp residues conserved in Motifs A and C (red asterisks in Figure 2A). Additionally, these 28 RTZ_LINEs had residues conserved in Motifs B’ and C (black asterisks in Figure 2A). These 28 RTZ_LINEs had Lys but not Gly residues conserved in Motif D. Amino acid substitution of Gly in Motif D is often observed in RNA-dependent RNA polymerases in negative-strand RNA viruses (Poch et al., 1989). Therefore, we held out the possibility that the RTZ_LTRs with amino acid substitution of Gly in Motif D were still functional. In RTZ_LINE-61, Gly residues in Motif E were substituted. However, given the conservation of the 2 critical amino acids for holding a bivalent metal ion, and the lack of a functional test, we held out the possibility that RTZ_LTR-61 might be still functional. Taken together, these results suggest that the RT domains of these 28 *rtz_LINE*s are potentially functional.

**Figure 2.**
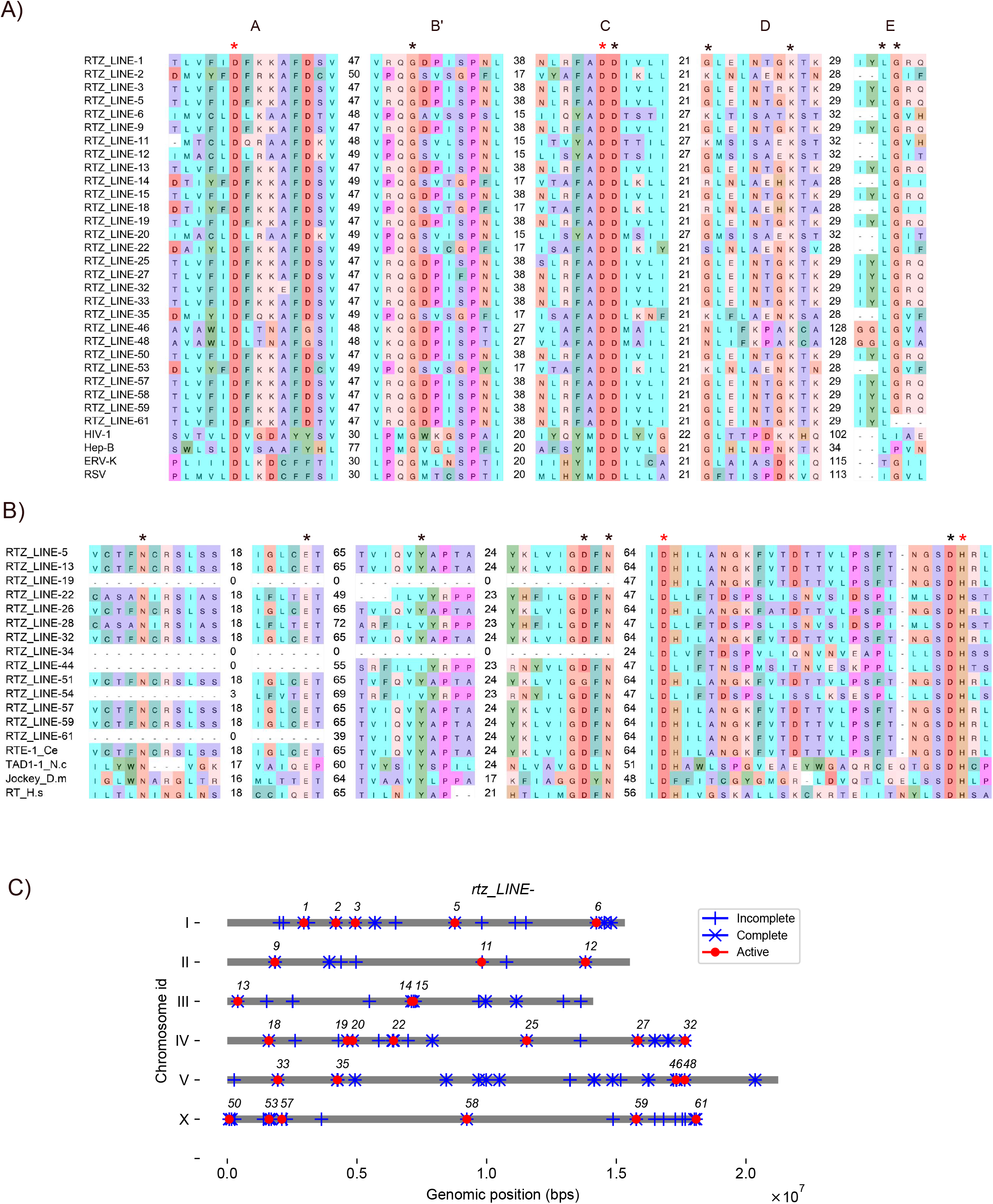
Potentially functional *rtz_LINE*s encoded in long interspersed nuclear elements (LINEs) (A) Alignment of reverse transcriptase (RT) domains of RTZ_LINEs with reference RTs. Abbreviations: HIV-1: Chain C, HIV-1 RT P66 subunit of human immunodeficiency virus type 1 [5TXO_C], Hep-B: hepatitis B virus RT_like family [QFR04538], ERV-K: Pol protein of human endogenous retrovirus K [CAA76882], RSV: Pol of Rous sarcoma virus [CAA48535]. (B) Alignment of the endonuclease (EN) domains of RTZ_LINEs with reference ENs. Abbreviations: RTE-1_Ce: apurinic-apyrimidic EN domain containing RT of non-LTR retrotransposon in *C. elegans* [AAC72298.1], AP_End_Hs: DNA-(apurinic or apyrimidinic site) EN in *H. sapiens* [NP_542379.1], TAD1-1_N.c: Exonuclease-Endonuclease-Phosphatase (EEP) domain containing Pol protein *N. crassa* [AAA21781.1], Jockey_D.m: EEP domain containing RT in *D. melanogaster* [AAA28675.1], RT_H.s: EN domain containing RT of L1, *H. sapiens* [AAB59368.1]. Asterisks indicate conserved residues. Red asterisks indicate residues for identifying potentially functional transposon-mobility genes. (C) Genomic positions of 28 potentially functional *rtz_LINE*s in *C. elegans* genome. Grey lines represent chromosomes. Red circles indicate positions of potentially functional genes of *rtz_LINE*-n. Numbers indicate numbers of *rtz_LINE* genes. Vertical ticks and × marks indicate incomplete and complete ORFs, respectively, of *rtz_LINE*s.

Next, by aligning the 61 RTZ_LINEs with reference ENs, we identified 14 RTZ_LINEs in which the critical Asp and His residues were conserved in their EN domains (red asterisks in Figure 2B). A previous experimental test of transposition activity in 138 human L1 copies revealed that the Asp and His residues are essential, but conserved amino acid residues in other motifs tolerate multiple mutations (Kines et al., 2016). Among the 14 RTZ_LINEs, RTZ_LINE-19 and -34 lacked a large N-terminal portion of the EN domain (Figure 2B). Therefore, we concluded that the remaining 12 RTZ_LINEs have potentially functional EN domains. Taken together, we concluded that the 6 *rtz_LINE*s encoded both potentially functional RT and EN domains (*rtz_LINE-5*, *rtz_LINE-13*, *rtz_LINE-22*, *rtz_LINE-57*, *rtz_LINE-59*, and *rtz_LINE-61*; Figures 2A and 2B and Supplementary Table 3).

Human L1 can retrotranspose via an EN-independent and RT-dependent mechanism (Morrish et al., 2002). Therefore, we concluded that the 28 *rtz_LINE*s with conserved RT domains (Figure 2A) are potentially functional genes for LINE transposition. These 28 functional *rtz_LINE*s were distributed across the entirety of each chromosome (Figure 2C).

### Transposon-mobility genes in TIRs in *C. elegans*

TIR is composed of 19 super families: hAT, Tc1/mariner, CACTA (En/Spm), Mutator (MuDR), P, PiggyBac, PIF/Harbinger, Mirage, Merlin, Transib, Novosib, Rehavkus, ISL2EU, Kolobok, Chapaev, Sola, Zator, Ginger, and Academ (Yuan and Wessler, 2011). The transposition of TIR elements is mediated by TP, which has a conserved DDD/E motif at the catalytic core (Yuan and Wessler, 2011). Structural analysis suggests that the DDD/E motif holds 2 metal ions to cleave and integrate the TIR dsDNA element (Davies et al., 2000; Lovell et al., 2002; Richardson et al., 2009). Mutation of the conserved DDD/E motif abolishes TP activity (Bolland and Kleckner, 1996; Naumann and Reznikoff, 2002). By aligning the 189 TPZs with reference TPs, we identified 94 TPZs that had DDD/E motifs conserved that were homologous to those of Tc1/mariner family TPs (Figure 3A and Supplementary Table 4). No TP aligned to DDD/E motifs of other TP super families. Thus, we considered these 94 *tpz*s to be potentially functional genes (Supplementary Table 4). The 94 functional *tpz*s were distributed across the entirety of each chromosome (Figure 3B).

**Figure 3.**
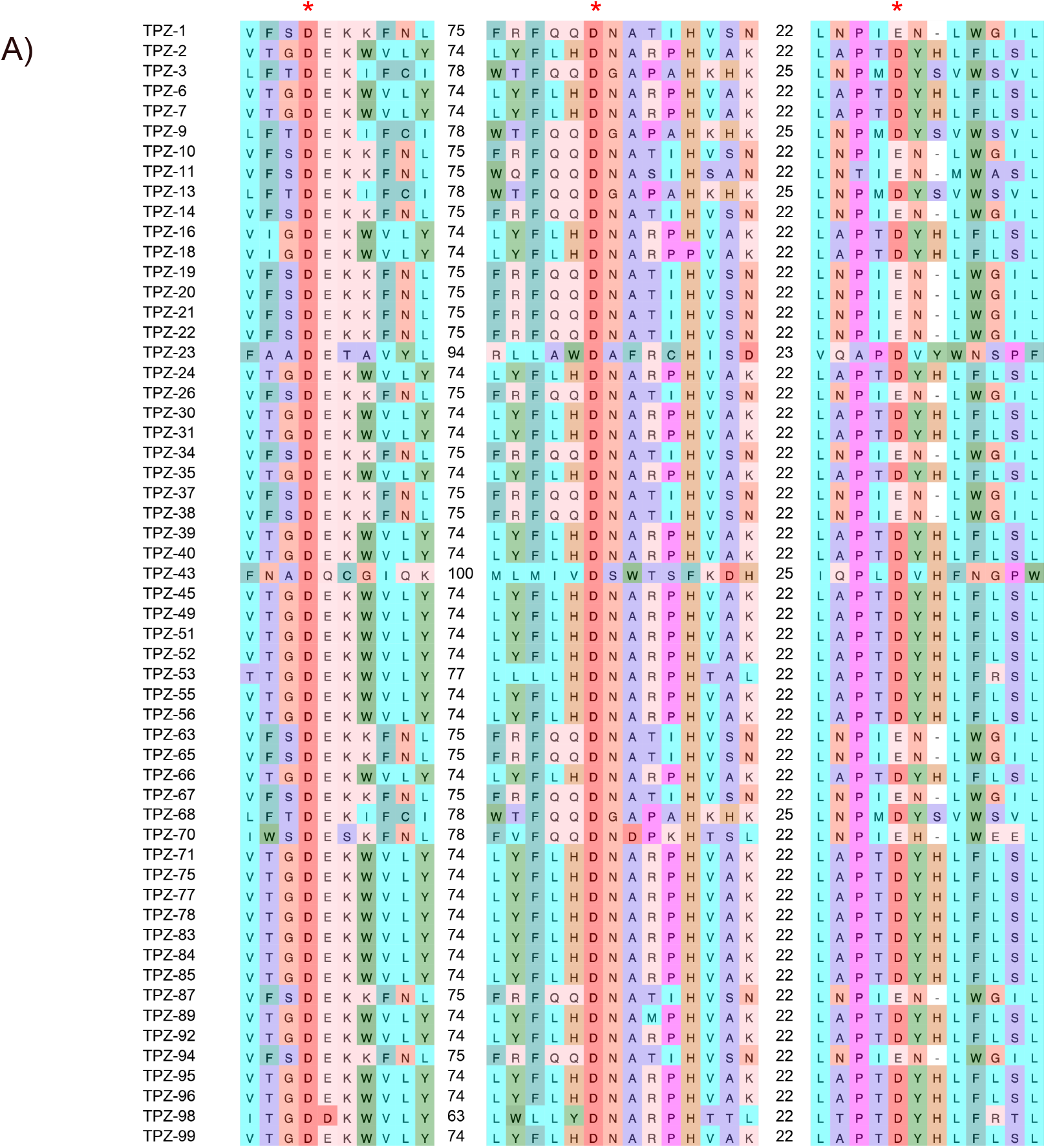
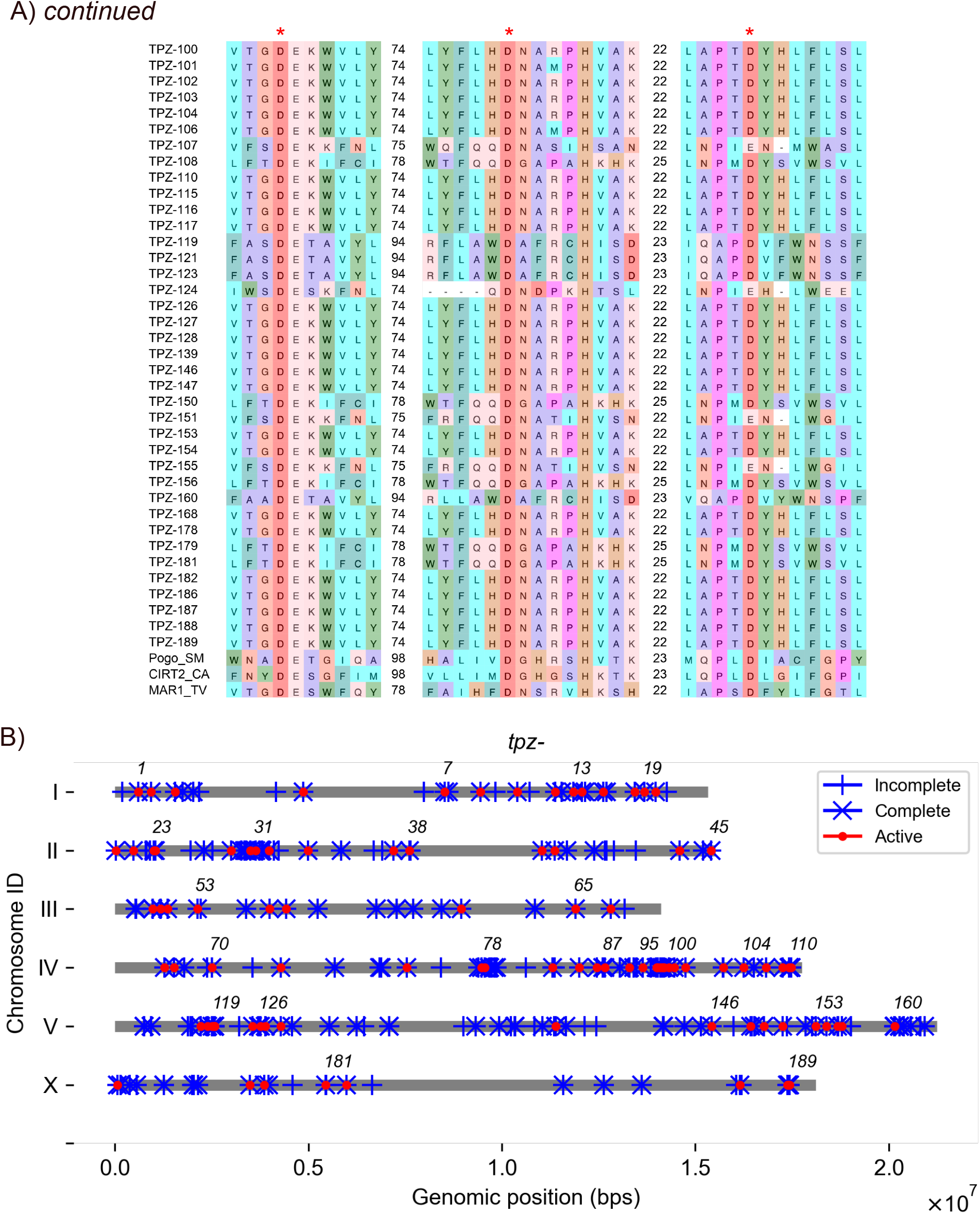
Potentially functional *tpz*s encoded in terminal inverted repeats (TIRs) (A) Alignment of catalytic core domains of TPZs with reference transposases (TPs). Amino acid sequences of Pogo_SM, CIRT2_CA, and Mar1_TV as examples of Tc1/mariner class TPs were arbitrarily obtained from (Yuan and Wessler, 2011). Red asterisks indicate conserved residues used to identify potentially functional transposon-mobility genes. (B) Genomic positions of 94 potentially functional *tpz*s in *C. elegans* genome. Grey lines represent chromosomes. Red circles indicate positions of potentially functional genes of *tpz*-n. Numbers indicate numbers of *tpz* genes. Vertical ticks and × marks indicate incomplete and complete ORFs, respectively, of *tpz*s.

### Transposon-mobility genes in Helitrons in *C. elegans*

Helitron transposition is mediated by a SF1 family helicase (HEL) with REP domain (REP-HEL). The 4 SF family HELs (SF1, SF2, SF3, and SF4) have conserved Motif I and Motif II domains. Motifs I and II correspond to the Walker A and Walker B domains, respectively, which are widely conserved among NTP-binding proteins (Hall and Matson, 1999; Walker et al., 1982). The Walker A/Motif I and Walker B/Motif II domains exhibit conservation of the Lys and Asp-Glu residues, respectively (Ambudkar et al., 2006; Gorbalenya and Koonin, 1993; Koonin, 1993; Walker et al., 1982). Crystallographic analysis showed that Lys in Motif I contacts with magnesium ion and β phosphate of ATP, and functions to stabilize the transition state during ATP hydrolysis. Asp-Glu residues in Motif II are also involved in ATP hydrolysis (Raney et al., 2013; Velankar et al., 1999). Mutations at Lys in Motif I or Asp-Glu in Motif II abolish HEL activity (Brosh and Matson, 1995; Graves-Woodward et al., 1997; Walker et al., 1997; Weng et al., 1996).

By aligning the 5 REP-HELs (RHZs) with reference HEL domains, we identified 5 RHZs that had Lys residues conserved in Motif I and had Asp-Glu residues conserved in Motif II (red asterisks in Figure 4A). Additionally, these 5 RHZs had: 1) weakly conserved residues in Motif Ia (black asterisks in Figure 4A), which functions for ssDNA binding and energy transfer from the ATP-binding site to the DNA-binding site (Raney et al., 2013); 2) conserved Gly-Asp and other resides in Motif III (black asterisk in Figure 4A), which is involved in contacting nucleotide γ-phosphates (Raney et al., 2013); 3) a conserved Arg residue in Motif IV, which may be involved in NTP hydrolysis (Raney et al., 2013); 4) conserved residues in Motif IV/V, the function of which is not well understood (Kapitonov and Jurka, 2001); 5) conserved residues in Motif V, which interacts with the sugar-phosphate backbone of DNA (Raney et al., 2013); and 6) conserved residues in Motif VI, which may form part of the ATP binding cleft to couple ATPase activity to HEL activity (Raney et al., 2013) (Figure 4A). Therefore, we considered these 5 *rhz*s to encode functional HEL domains.

**Figure 4.**
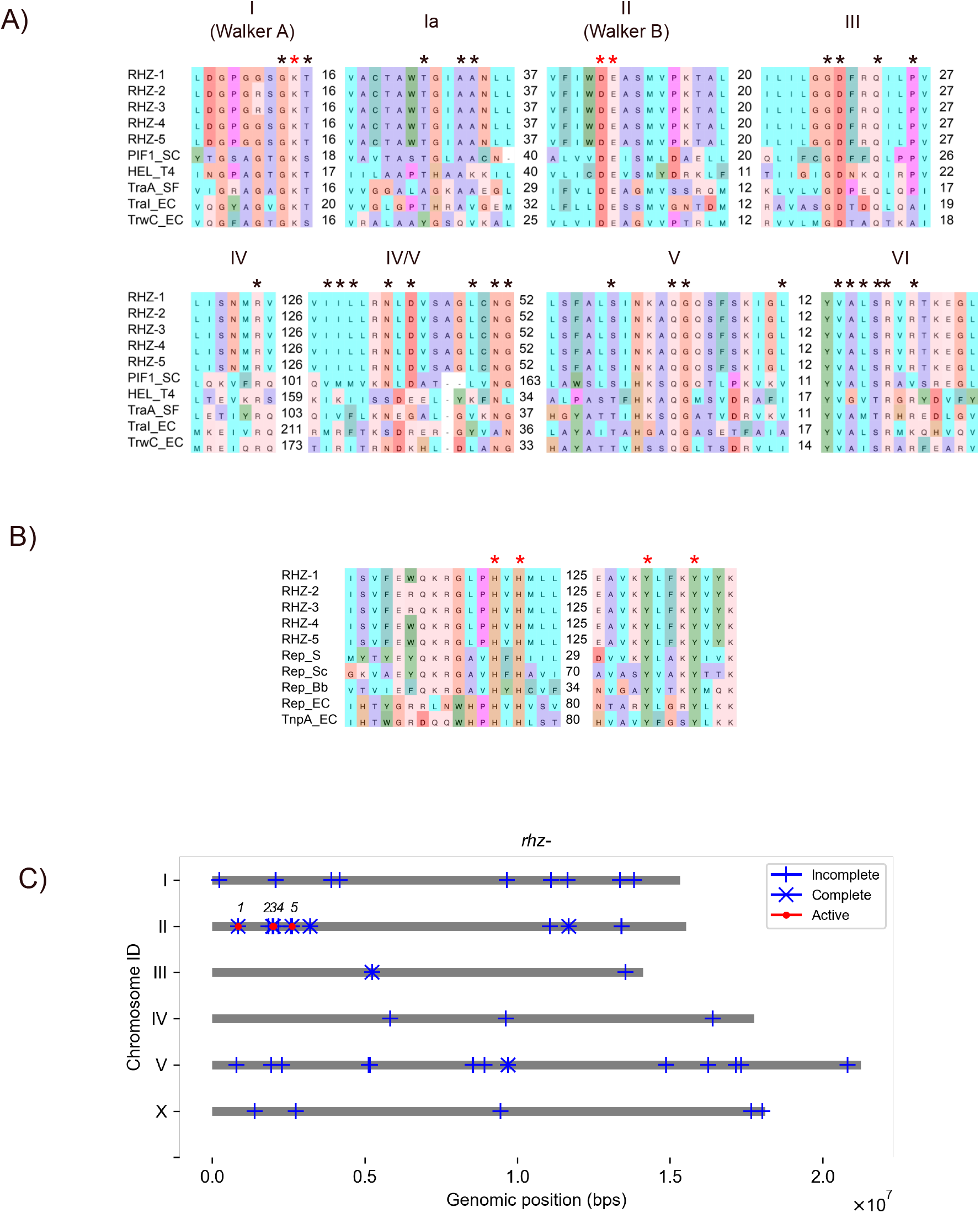
Potentially functional *rhz*s encoded in Helitrons. (A) Alignment of amino acid sequences of helicase (HEL) domains with reference HELs. PIF1 in *S. cerevisiae* [P07271], HEL_T4 in Enterobacteria phage T4 [P32270], TraA *S. fredii* [P55418], TraI_EC *E. coli* [P14565], TRWC *E. coli* [Q47673]. (B) Alignment of amino acid sequences of REP domains with reference REPs. Rep_Bb in *B. borstelensis* [BAA07788.1], Rep plasmid pVT736-1 [AAC37125.1], Rep_Pf3 P. phage Pf3 [AAA88392.1], Rep_EC IS91 [BCN22733.1], TnpA_EC IS91 TnpA *E. coli* [QIC00531.1]. Asterisks indicate conserved residues. Red asterisks indicate residues used for identifying potentially functional transposon-mobility genes. (C) Genomic positions of 5 potentially functional *rhz*s in *C. elegans* genome. Grey lines represent chromosomes. Red circles indicate positions of potentially functional genes of *rhz*-n. Numbers indicate numbers of *rhz* genes. Vertical ticks and × marks indicate incomplete and complete ORFs, respectively, of *rhz*s.

In the REP domain, the HUH Y2 motif (in which U is a hydrophobic residue) is evolutionarily conserved. HUH holds divalent metal ions to form nicks in the DNA strand in EN activity, whereas Tyr residues form a transient covalent bond with the cleaved DNA strand to generate phospho-tyrosine for DNA strand transfer. The EN activity is abolished by mutation of either 2 His or 2 Tyr residues (Grabundzija et al., 2016). By aligning the 5 RHZs with the reference REP domains, we found that the 5 RHZs conserved the 2 His and 2 Tyr residues (red asterisks in Figure 4B). Taken together, our results suggest that the 5 *rhz*s encode both potentially functional HEL and EN domains (Supplementary Table 5). These 5 *rhz*s are located only on chromosome II (Figure 4C).

### Transposon-mobility genes in Maverick/Polinton (MP) elements in *C. elegans*

Our RepeatMasker (RM) analysis did not identify MP elements. MPs are reportedly located on chromosomes I, II, III, IV, and X of the *C. elegans* genome (Feschotte and Pritham, 2005; Gao and Voytas, 2005; Pritham et al., 2007). Using the representative AL110478.1 sequence of MP in *C. elegans* (Pritham et al., 2007), we searched for MP copies in VC2010 by using the NCBI nucleotide BLAST program (https://blast.ncbi.nlm.nih.gov/). We identified short homologous regions >100 bps that were densely scattered on 3 distinct regions on chromosome I (MP Ia, MP Ib, and MP Ic), and 2 regions on chromosome X (MP Xa and MP Xb) (Figure 5A and 5B). Thus, the reference MP copy used in our homology search was aligned discontinuously in these MP copies in the VC2010 genome assembly (Figure 5B). Additionally, MP copies were located on different chromosomes from previous reports.

**Figure 5.**
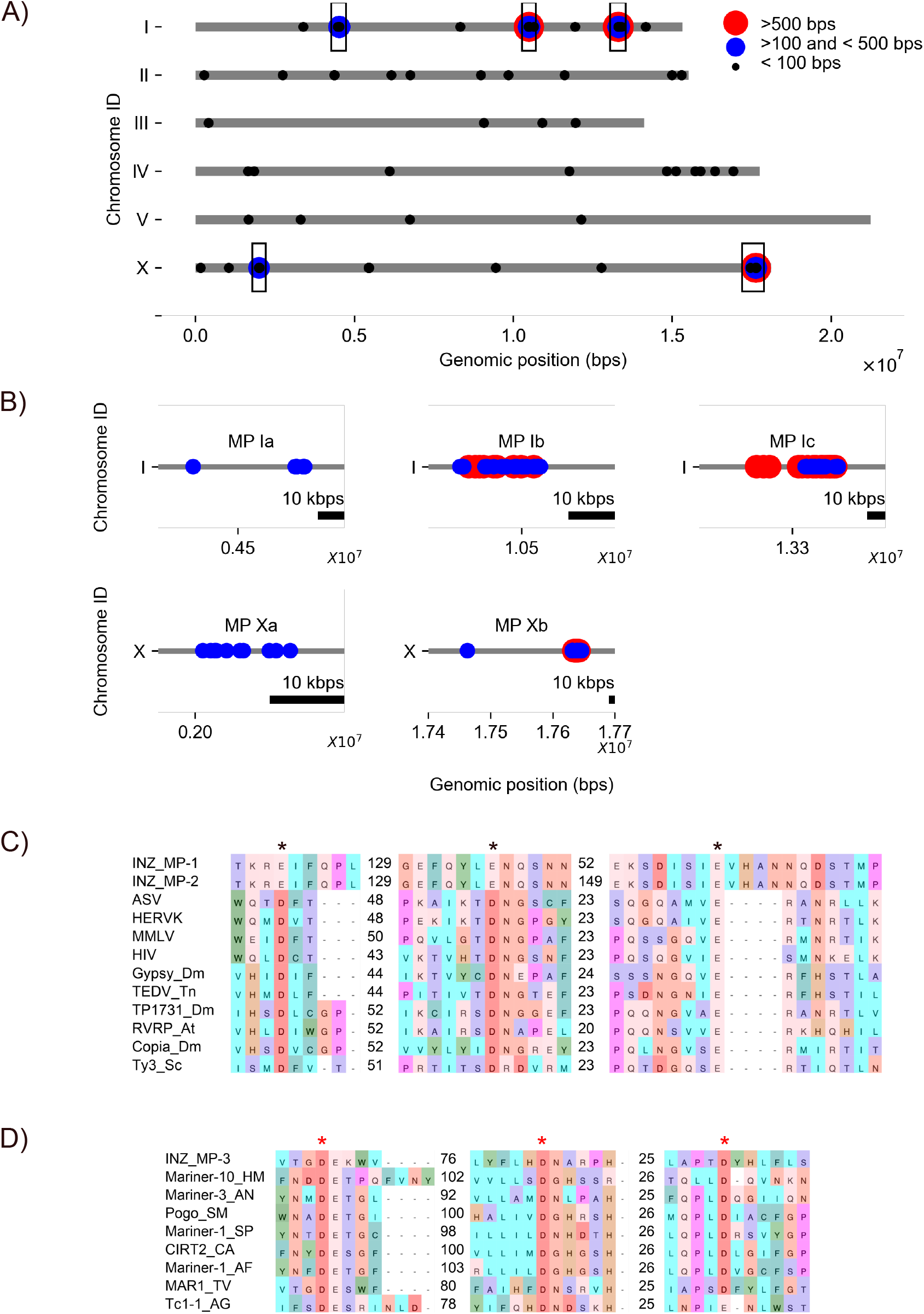
Maverick/Polinton (MP) elements in *C. elegans* genome. (A) Genomic positions of homologous regions with AL110478.1. Five rectangular regions are as follows: in chromosome I: MP Ia, 3543-bp region from 0.4521550×10^7^ to 0.4525093×10^7^ bps; MP Ib, 17,276-bp region from 1.0486778×10^7^ bps to 1.0504054×10^7^ bps, and MP Ic, 43,817-bp region from 1.3280931×10^7^ bps to 1.3324748×10^7^ bps; and in chromosome X: MP Xa, 11,758-bp region from 0.2000999×10^7^ bps to 0.2012757×10^7^ bps, and MP Xb, 183,668-bp region from 1.7462913×10^7^ bps to 1.7646581×10^7^ bps. (B) Scattered distribution of homologous DNA regions with AL110478.1 in each of 5 MP copies. Red dots indicate homologous regions >500 bps. Blue dots indicate homologous regions of <500BPS and >100 bps. (C) Alignment of catalytic core domains of INZ_MP-1 and INZ_MP-2 with reference integrases (INs). Abbreviations: ASV: IN in avian sarcoma virus [1ASU_A], HERVK: Pol protein in human endogenous retrovirus K [CAA76885], MMLV: p46 IN Moloney murine leukemia virus (MoMLV) [NP_955592.1], HIV: IN in HIV-1 [1BIZ_A], Gypsy_Dm: IN, Gypsy endogenous retrovirus in *D. melanogaster* [CAB69645], TEDV_Tm: ORFB in TED virus in *T. ni* [YP_009507248], TP1731: Pol polyprotein in transposon_1731 in *D. melanogaster* [S00954], RVRP_At: retrovirus-related like Polyprotein in *A. thaliana* [CAB78488_1], Copia_Dm: Gag-Int-Pol protein in COPIA in *D. melanogaster* [P04146], Ty-3_Sc: Gag-Pol polyprotein in Ty3-G in *S. cerevisiae* [GFP69998.1]. (D) Alignment of catalytic core domains of INZ_MP-3 with reference transposases (TPs). Amino acid sequences of Mariner-10_HM, Mariner-3_AN, Pogo_SM, Mariner-1_SP, CIRT2_CA, Mariner-1_AF, MAR1_TV, Tc1-1_AG as examples of Tc1/mariner class TPs were arbitrarily obtained from (Yuan and Wessler, 2011). Asterisks indicate conserved amino acids in catalytic core.

The VC2010 genome assembly provides substantial advantages over its predecessors in both precision and completeness, but at the current stage, any assembly is imperfect (Yoshimura et al., 2019). Compared to the N2 genome assembly, the VC2010 assembly contains short insertions, deletions, and duplications ranging from tens to thousands of base pairs, which are distributed in all chromosomes (Yoshimura et al., 2019). These differences could be due to polymorphism in the PD1074 strain that was used to obtain the VC2010 assembly, or could be due to errors arising during sequencing and/or assembly of the N2 genome assembly. Therefore, the discontinuity within the MP copies (Figure 5A and 5B) is likely to be the actual sequence present in the PD1074 strain. On the other hand, artifactual large structural variations in VC2010, as we found in chromosomal locations of the MP copies, have been suggested to remain even after careful correction in the VC2010 assembly (Yoshimura et al., 2019). Thus, it remains inconclusive whether the different chromosomal locations of the MP copies in VC2010 are actually present in the PD1074 strain or due to errors in the assembly process to obtain the VC2010 assembly.

To identify transposon-mobility genes encoded in these MP copies, *i.e.*, DNA polymerase B (DNA POLB) and IN, we applied SNAP and DIAMOND to the 5 MP copies. Two DNA POLB-related genes (*polB_MP*) encoding 378 and 93 amino acids were located in MP Ia and MP X, respectively. Two IN genes (*inz_MP*) were located in MP Ib and MP Ic. Interestingly, a gene encoding the helix-turn-helix 28 (HTH28) domain-containing protein, which is conserved in some TPs (shttps://www.uniprot.org/uniprotkb?query=HTH28%20transposase), is located in MP Xa. Because TPs are functionally and evolutionarily related to INs (Capy et al., 1996, 1997), we considered these 3 IN-related genes to be *inz_MP*s.

DNA POLB has 5 conserved motifs from Motifs I to V (Iwai et al., 2000). By aligning PolB_MP-1 (at MP Ib) and PolB_MP-2 (at MP Xa) with the reference DNA POLBs, we found that PolB_MP-1 lacked most of the N-terminal motifs from I to III, and only had YnDTD conserved in Motif IV. Additionally, PolB_MP-2 only had conservation of a short N-terminus fragment, and did not exhibit conservation of Motifs I to V. Therefore, we considered that these PolB_MPs do not encode functional proteins. Next, by aligning INZ_MP-1 (at MP Ib), -2 (at MP Ic), and -3 (MP Xa) with reference INs, we found that INZ_MP-1 and -2 aligned their EEEs with the reference DDE triad (black asterisks in Figure 5C). Asp and Glu have the same negatively charged carboxylate groups. We conclude that *inz_MP-*1 and -2 are potentially functional genes. INZ_MP-3 did not align with the reference DDE triads, whereas INZ_MP-3 aligned with the DDD/E triad in the Tc1 family TPs (red asterisks in Figure 5C). We also considered that *inz_MP-3* to be a potentially functional gene.

The IN encoded in a MP copy has been identified as a cellular IN (c-integrase) that is homologous to the retrotransposon IN (Gao and Voytas, 2005). Additionally, the IN in MP is highly homologous to the TP encoded in ginger DNA transposon (Bao et al., 2010). MP has been proposed to evolutionarily play a role as a hotbed that exchanges genes between eukaryotic DNA mobile elements (dsDNA viruses, adenoviruses, small ssDNA viruses, Mavirus-like virophages, icosahedral viruses) (Koonin et al., 2015; Krupovic and Koonin, 2015; Yutin et al., 2015). Considering that *inz_MP-3* at MP Xa encodes the HTH28 domain, and that the DDD motif is homologous to the Tc family of DNA TPs, we conclude that the Tc family of DNA transposons may be involved in MP evolution.

### Transposon-mobility genes in DIRS, PLE, and Crypton elements in *C. elegans*

Our RM analysis did not identify copies of DIRS or Crypton. A previous homology-based search of 34 nematode species to identify Tyr-REC genes that mediate transposition of DIRS and Crypton identified an incomplete cDNA for a Tyr-REC gene in chromosome II of *C. elegans* (Szitenberg et al., 2014). We applied SNAP to a genomic region of this incomplete cDNA with 20-kbp 5’ flanking and 20-kbp 3’ flanking genomic regions in VC2010, but did not find any complete ORF. From this result and previous studies, we conclude that *C. elegans* may not have a functional Tyr-REC gene. Finally, consistent with a previous report (Arkhipova, 2006), our analysis on VC2010 did not identify any PLE copies.

## Discussion

Owing to the development of the user-friendly algorithm RM, the genomic ratio and number of transposon copies have been studied comprehensively across all transposon orders in various species (Wells and Feschotte, 2020; Wicker et al., 2007). An ORF structural analysis of RT genes encoded in LINE contributed to an estimation of autonomously mobile copies of LINE across various species (Ivancevic et al., 2016). Additionally, critical amino acids at the catalytic core of transposon-mobility genes have been studied in LTRs (Bowen and McDonald, 1999; Ganko et al., 2001; Kanzaki et al., 2018), LINEs (Bessereau, 2006; Marín et al., 1998; Youngman et al., 1996), and TIRs (Fischer et al., 2003) in *C. elegans*. In this report, we comprehensively studied transposon-mobility genes in all transposon orders in the latest genome assembly of *C. elegans* (VC2010). According to our RM analysis, more than 60,000 transposon copies exist in the latest *C. elegans* genome assembly (Figure 6A and Table 1). These transposon copies encode 428 complete ORFs, 66.8% of which (285 genes) are transposon-mobility genes. Among them, we identified 142 potentially functional genes, including 8 *rtz_LTR*s, 7 *inz_LTR*s, 28 *rtz_LINE*s, 94 *tpz*s, and 5 *rhz*s (Figure 6B). Additionally, our manual analysis identified 3 potentially functional *inz_MP*s (Figure 6B). In total, we identified 145 potentially functional transposon-mobility genes. Thus, in contrast to the large number of transposon copies in *C. elegans*, only a limited number are autonomously mobile. Because we did not analyze DNA elements encoding functional protein domains other than those studied in this report or that are required for protein expression, the number of transposon-mobility genes that are truly functional *in vivo* is likely to be fewer than 145.

**Figure 6.**
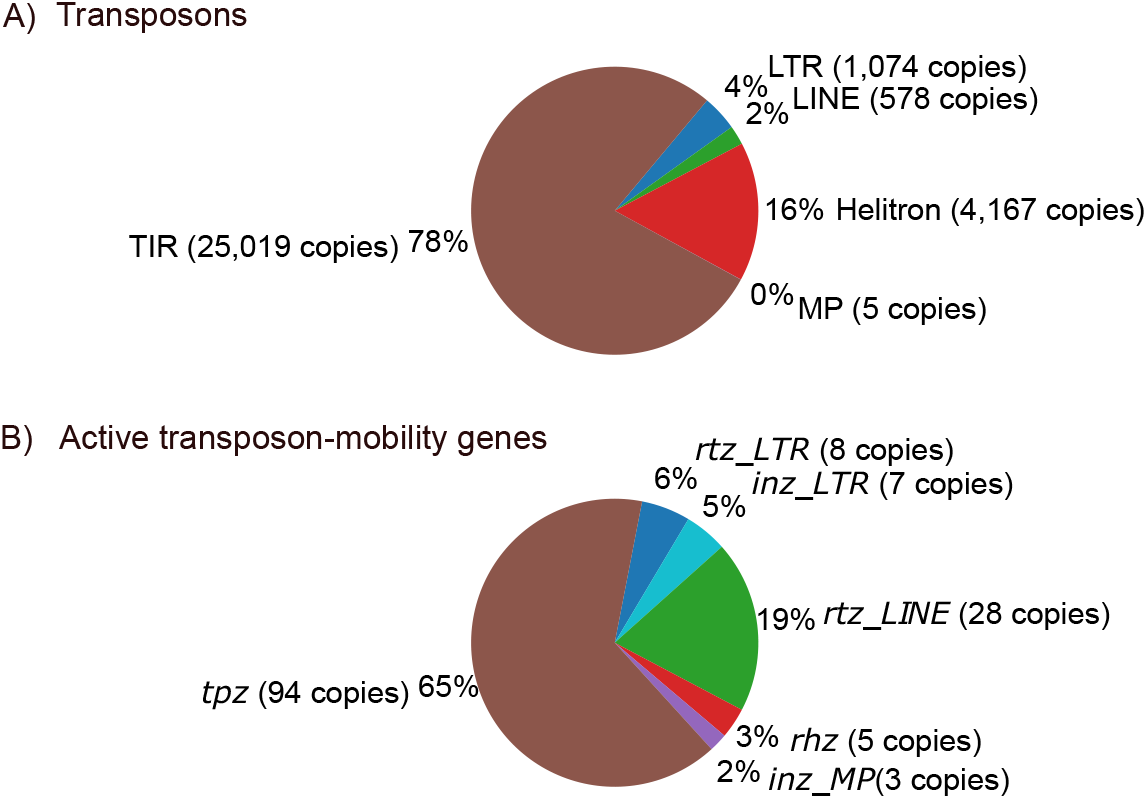
Summary of potentially functional transposon-mobility genes in *C. elegans* genome. (A) The percentage of copy numbers of LTR (blue), LINE (green), Helitron (red), MP (purple), and TIR (brown) transposons in the total copy number of transposons in *C. elegans* genome. (B) The percentage of potentially functional transposon-mobility genes; 8 *rtz_LTR* (blue) and 7 *inz_LTR* (light blue), 28 *rtz_LINE* (green), 5 *rhz* (red), 3 *inz_MP* (purple), and 94 *tpz* (brown) in total 145 potentially functional transposon-mobility genes identified in this study.

We compared the 145 potentially functional transposon-mobility genes identified in this study to those identified previously. In previous studies of LTR retrotransposons, researchers identified 24 (Ganko et al., 2001) or 10 (Kanzaki et al., 2018) full-length copies among 124 or 62 LTR copies, respectively, in *C. elegans*. Another study reported 17 *rtz_LTR*s with conserved Asp residues in Motifs A and C, and 15 *inz_LTR*s with a conserved DDE triad (Bowen and McDonald, 1999). We identified 8 potentially functional *rtz_LTR* and 7 potentially functional *inz_LTR* genes. Thus, we found fewer potentially functional *rtz_LTR*s and *inz_LTR*s in this study than were identified by previous studies (Bowen and McDonald, 1999; Ganko et al., 2001; Kanzaki et al., 2018). Differences in the genome assembly used in (Bowen and McDonald, 1999; Ganko et al., 2001), as discussed in (Yoshimura et al., 2019), might contribute to this discrepancy. On the other hand, using the same VC2010 assembly as us, Kanzaki et al. (2018) identified 10 LTR copies (Kanzaki et al., 2018); we identified nearly the same number but slightly fewer transposon-mobility genes. A detailed analysis of the transposon-mobility genes in the coding sequence of the catalytic core of enzymes in the 10 full-length LTR copies (Kanzaki et al., 2018) may provide a more consistent number of potentially functional LTR genes in our study. Overall, why we obtained fewer LTR copies in our study remains unclear. Nevertheless, transposon-mobility genes in the most-studied LTR, Cer1 on chromosome III (Ganko et al., 2001), were found on our list of potentially functional genes as *rtz_LTR-5* and *inz_LTR-5* (Figure 1C). Cer1 is both biologically active (Dennis et al., 2012; Moore et al., 2021; Sun et al., 2023) and mobile in recent evolutionary history, based on a comparison of natural isolates of *C. elegans* (Palopoli et al., 2008).

For LINE retrotransposons, over 1,000 copies of LINE (Bessereau, 2006), 6 copies of the RTE LINE suborder (Youngman et al., 1996), and 17 copies of the T1/CR1 LINE suborder (Marín et al., 1998) encode the *rtz_LINE* that conserves the Asp residues in Motifs A and C. We identified a similar number of LINE copies (618 copies; Table 1). Among them, 28 *rtz_LINE*s preserved potentially functional RT domains, which is more than total number of potentially functional *rtz_LINE*s reported previously. For TIR DNA transposons, a previous report showed that 61 *tpz*s had the conserved DDD/E motifs in 127 copies of Tc/mariner family TIR (Fischer et al., 2003). We identified 94 potentially functional *tpz*s in 189 copies of the Tc/mariner family TIR, a larger number than was reported previously. For Helitron DNA transposon, we found that 1.65% of the *C. elegans* genome was occupied with Helitron copies (Table 1), similar to previous reports (~2%) (Kapitonov and Jurka, 2001). There was no further amino acid sequence analysis about *rhz*s. Thus, the potentially functional transposon-mobility genes identified in this study among all transposon orders include most of the previously identified genes, as well as a set of new genes.

In human and mouse genomes, L1 is the most abundant among transposon classes, with 145 copies in human and 2,811 copies in *Mus musculus* being conserved at full length (Lander et al., 2001; Penzkofer et al., 2017; Waterston et al., 2002). In *Drosophila melanogaster*, *Saccharomyces cerevisiae*, and *Arabidopsis thaliana* (Zhang and Wessler, 2004), LTR is the most abundant class, with 325 copies in *D. melanogaster* (Bergman et al., 2006) and 51 copies in *S. cerevisiae* (Kim et al., 1998) being conserved at full length. LTR is only the transposon encoded in *S. pombe*, with 13 copies being conserved at full length (Bowen et al., 2003). DNA transposons are the most abundant in *Danio rerio* (zebrafish), with 2.3 million copies, but it is unknown how many of these copies are active (Howe et al., 2013). Similarly, it is unknown how many of the 286 LTR copies in *A. thaliana* are active (Zhang and Wessler, 2004). In our study, TIR was the most abundant transposon class (20,852 copies; Table 1) in *C. elegans*, encoding 94 copies of potentially functional *tpz*. Our identification of 145 potentially functional transposon-mobility genes indicates that

*C. elegans* encodes the fewest number of potentially functional transposon-mobility genes among conventional metazoan model organisms. We propose that *C. elegans* can be a useful model metazoan to study the conserved physiological, pathological, and evolutionary roles of transposon-mobility genes.

## Methods

### Bioinformatics analysis

VC2010 was downloaded from https://www.ebi.ac.uk/ena/browser/view/UNSB01. To identify transposons, RepeatMasker (RM) version 4.1.0 (http://www.repeatmasker.org/) was used with options -no_is for skipping bacterial insertion element check, -s: slow search for more sensitivity, and -pa 8 for sequencing batch jobs to run in parallel. For RM analysis, we used a library of *C. elegans* transposon sequences, which was isolated from RepBase RepeatMasker libraries (Bao et al., 2015) (Supplementary Material 1). To identify ORFs, SNAP version 2006-07-28 (Korf, 2004) was used. To infer the function of proteins encoded in ORFs, DIAMOND (v2.0.2.140) (Buchfink et al., 2014) was used. UniRef50 protein ID produced by DIAMOND was translated into functional protein IDs using the website Uniprot (https://www.uniprot.org/uploadlists/). To align amino acid sequences, MAFFT v7.453 (Katoh and Standley, 2013; Katoh et al., 2002) was used. A sequence alignment viewer was downloaded from https://github.com/dmnfarrell/teaching/blob/master/pyviz/bokeh_sequence_align.ipynb.

## Acknowledgements

The authors thank Dr. Damien Farrell at University College Dublin, US for the sequence alignment viewer. We thank WormBase (www.wormbase.org) for a valuable help for our analysis. This study was supported by JSPS KAKENHI grant number 20K20321 (Grant-in-Aid for Challenging Research (Exploratory))(Y.A.).

